# Testing the link between isoaspartate and Alzheimer’s disease etiology

**DOI:** 10.1101/2022.05.03.490418

**Authors:** Jijing Wang, Cong Guo, Zhaowei Meng, Marissa D. Zwan, Xin Chen, Sven Seelow, Susanna L. Lundström, Sergey Rodin, Charlotte E. Teunissen, Roman A. Zubarev

## Abstract

Isoaspartate (isoAsp) is a damaging amino acid residue formed in proteins as a result of spontaneous deamidation. IsoAsp disrupts the secondary and higher order structures of proteins, damaging their functions and making them prone to aggregation. An association has been suggested between isoAsp and Alzheimer’s Disease (AD). Here we strengthened the link between isoAsp and AD by novel approaches to isoAsp analysis in blood human serum albumin (HSA), the most abundant blood protein, a major carrier of amyloid beta (Aß) peptide and phosphorylated tau (pTau) protein in blood and a key participant in their clearance pathway. We discovered a reduced amount of anti-isoAsp antibodies (P < .0001), an elevated isoAsp level in HSA (P < .001), more HSA aggregates (P < .0001) and increased levels of free Aß (P < .01) in AD blood compared to healthy controls. We also found that deamidation significantly reduces HSA capacity to bind with Aß and pTau (P < .05). These findings support the presence in AD of a bottleneck in clearance of Aß and pTau leading to their increased concentrations in brain and facilitating their aggregations there.

**RESEARCH IN CONTEXT:**

1. **Systematic review:** We reviewed the evidence that associates isoaspartate (isoAsp) residue in blood proteins with the etiology of Alzheimer’s disease (AD). However, the link between isoAsp in blood and aggregation of amyloid beta (Aß) peptide and phosphorylated tau (pTau) protein in brain remained unclear.
2. **Interpretation:** For the first time we demonstrate that isoAsp-containing human serum albumin (HSA) forms aggregates with reduced binding capacity toward Aß peptide and pTau protein. Using a novel ELISA, we discovered in AD blood elevated levels of isoAsp in HSA, together with reduced endogenous anti-isoAsp antibody levels, suggesting hampered Aß and pTau clearance in AD.
3. **Future directions:** As degradation of the innate anti-isoAsp defenses may take years to develop, investigation of the isoAsp role in early stages of AD is warranted. And enrollment of different neurodegenerative disease cohorts will illustrate if isoAsp is AD-specific or universal to diseases related to aging.

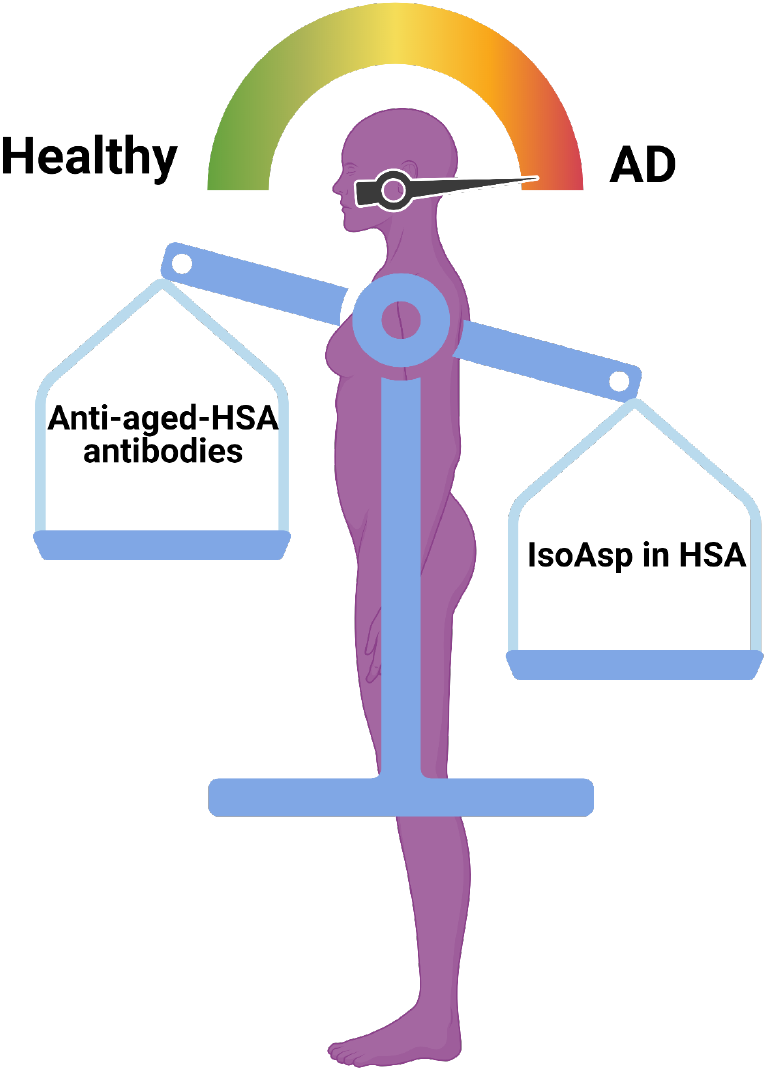

## 1. NARRATIVE

### 1.1 Contextual background

Alzheimer’s disease (AD) is the most common form of senile dementia, for which there is currently no cure. The onset of AD is traditionally attributed to formation in brain of amyloid plaques due to accretion of amyloid-beta (Aβ) peptides and tangles formed by aggregating hyper-phosphorylated Tau (pTau) protein [1]. Despite the fact that over 99% of the clinical trials based on Aβ hypothesis have failed, including several recent attempts [2], formation of Aß plaques and pTau tangles remains the strongest hallmark of AD, and the only FDA approved disease modifying drug [3]. The recently gained ability to measure Aß and pTau levels in blood for AD diagnostics using sophisticated antibody approaches has generated much enthusiasm in the AD community [4–7]. Therefore, it is beneficial for any AD etiology hypothesis to explain accumulation of Aß and pTau in brain, even though the true cause of AD may lie elsewhere. One of the less exploited AD origin theories can be traced back to the protein aging hypothesis proposed in the 1990s, suggesting that the accumulation of isoaspartate (isoAsp) residues resulting from deamidation of asparaginyl (Asn) and isomerization of aspartyl (Asp) residues triggers Aβ aggregation [8–10]. Although the link between isoAsp and pTau has not been established, this hypothesis deserves thorough consideration.

Deamidation of Asn is by far the most abundant posttranslational modification in proteins [11]. It is a spontaneous chemical reaction proceeding at a high rate even at physiological conditions [12]. As soon as the protein molecule is expressed, deamidation starts. Ammonia loss from Asn results in cyclic succinimide (Succ) (Figure 1A), while an alternative and less frequent pathway to Succ is the water loss from the L-Asp residue. Succinimide residue is unstable and easily hydrolyzes by attaching a water molecule, which most often (in 70-75% of the cases) results in isoAsp, a ß-isomer of the Asp residue [11].

**Figure 1.**
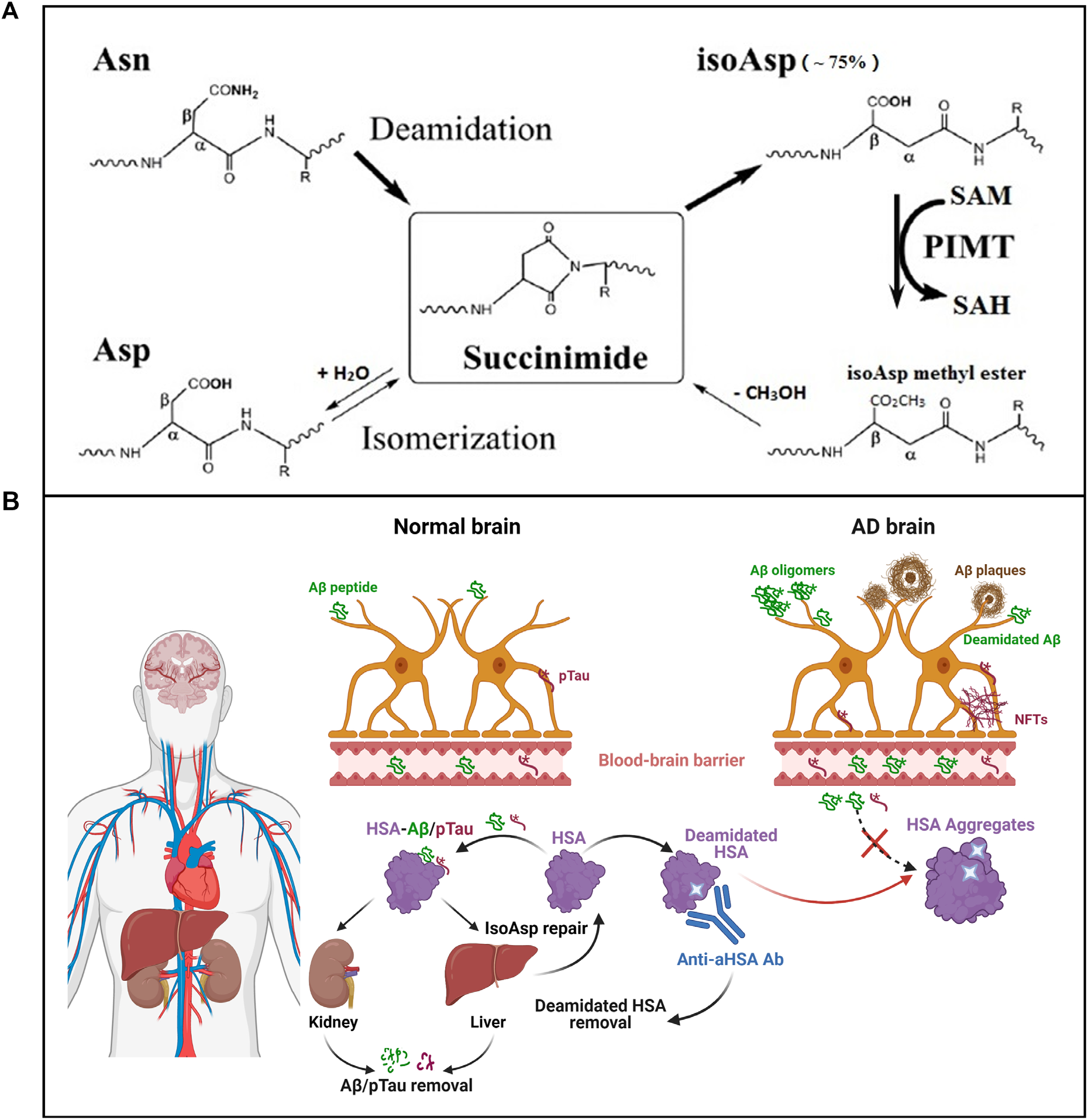
IsoAsp formation and the updated isoAsp hypothesis of AD. (A) Formation of isoAsp via succinamide intermediate by deamidation of Asn or isomerization of Asp residues, and its “repair” by the enzyme PIMT that methylates isoAsp using SAM as a methyl donor, converting it to SAH. Methylated isoAsp spontaneously loses methanol molecule, to become either normal L-Asp (in ≤25% cases) or isoAsp again. (B) The updated isoAsp hypothesis of AD: Aß peptide and pTau protein produced in brain are cleared by passing via blood brain barrier and being carried by HSA to kidneys and liver. Spontaneously deamidated HSA is repaired in liver; failure of repair leads to HSA aggregation, with a lost ability to bind Aβ and pTau. The anti-isoAsp antibodies remove deamidated HSA, and newly synthesized HSA molecules restore homeostasis. Due to a combination of insufficient repair and reduced removal of deamidated HSA, the diminished clearance of Aβ and pTau causes their accumulations in brain, with the Aβ oligomers and ultimately plaques as well as neurofibrillary tangles formed, which results in AD. Image was created via BioRender. Abbreviations: isoAsp, isoaspartate; Asn, asparagine; Asp, aspartate; PIMT, L-isoaspartyl methyltransferase; SAM, S-adenosylmethionine; SAH, S-adenosylhomocysteine; AD, Alzheimer’s disease; HSA, human serum albumin; Aβ, amyloid beta peptide; pTau: phosphorylated Tau; NFTs: neurofibrillary tangles; Ab: antibody.

In isoAsp, a CH2 group is rearranged from the side chain and inserted between the alpha-carbon and carbonyl group, which extends the polypeptide backbone. This extension disrupts the protein secondary structure, such as α-helices and ß-sheets, and often hampers protein function. As an example, deamidated human epidermal growth factor (EGF) no longer binds to its receptor, and thus loses its activity [13]. With most proteins folded into a native structure that is hydrophobic inside, isoAsp-induced structural changes often reduce protein solubility, making protein molecules prone to aggregation. In addition, isoaspartyl bonds are resistant to most proteolytic enzymes [14, 15] and can be cleaved only after the protein is degraded to a dipeptide by a few known isoaspartyl dipeptidases [16–18]. Difficulties to degrade isoaspartyl peptides proteolytically in antigen-presenting cells may render such peptides immunogenic [19].

Consistent with the link between isoAsp build-up and AD, we have earlier found elevated isoAsp levels in abundant blood proteins at all stages of AD compared to healthy controls [20]. Later, we confirmed that the blood proteins associated with rapid AD development are easily aggregated proteins, such as fibrin and fibrinogen [21]. These proteins tend to interact with amyloid peptides, contributing to their fibrillization and the formation of degradation-resistant fibrin clots in brain [21, 22]. Protease inhibitors have also been found correlating with AD, while the proteins delaying the AD development were those associated with clearance of protein aggregates, such as proteases, as well as immune-response proteins [21]. Both studies have been conducted by a combination of liquid chromatography and mass spectrometry (LC-MS/MS) – a comprehensive but expensive method of analysis. The isoAsp content in abundant blood proteins as well as the levels of these proteins could each provide an ≈ 80% accuracy in separating AD and healthy controls [20], which supported the isoAsp hypothesis but left room for a hope that more accurate type of analysis would be more predictive.

Although deamidation of asparaginyl is considered irreversible, there is an enzyme protein L-isoaspartyl methyltransferase (PIMT) that mitigates its damaging effect. PIMT methylates isoAsp using as a methyl donor S-adenosylmethionine (SAM) that becomes S-adenosylhomocysteine (SAH). Methylated isoAsp spontaneously loses methanol molecule to become in 16-25% cases normal L-aspartate (L-Asp), while with an overwhelming probability methyl loss (or methanol loss followed by water attachment) yields isoAsp again [23]. This low-efficiency quasi-repair (yielding L-Asp instead of the original L-Asn) occurs mostly in the liver, where the greatest majority of SAM is produced and consumed, and which the blood flow passes 20–25 times per day. The most abundant protein in blood, human serum albumin (HSA), is also produced mostly in the liver. HSA circulates in blood for 25 days on average, thus passing through the liver 500-600 times before removal. On each passage, PIMT and SAM convert the isoAsp residues to L-Asp, largely restoring protein structure and solubility. However, with age SAM production declines [24], rendering repair mechanism insufficient, which may result in isoAsp accumulation in long-lived proteins, including HSA.

HSA plays a special role in AD etiology. Being a major carrier protein, HSA is one of the most potent Aβ sequestering agents, binding 90% to 95% of the Aβ in blood plasma [25]. Model organism studies indicated that serum albumin inhibits Aβ aggregation [26, 27] and fibrillation [28–30] by binding Aβ monomers and oligomers, while human studies have revealed the association of low HSA levels with cognitive impairment [31, 32] as well as AD [33, 34]. A recent rather large study (n = 396) demonstrated that the HSA level is inversely associated as a continuous variable with Aβ deposition and Aβ positivity in older adults [35]. This result is in agreement with the earlier finding that HSA can regulate Aß peptide fiber growth in the brain interstitium [29]. Not much is known about binding of pTau to HSA, but infusing HSA intracerebroventricularly in an AD 3xTg mice model reduced both total Tau and pTau [36].

The Aß peptide is produced in the brain by astrocytes and neurons and secreted into the extracellular space, where it can meet one of the two basic fates. One fate is the clearance by enzymatic degradation or receptor-mediated export to blood plasma or cerebrospinal fluid. Not enough is known of Aβ clearance pathway out of the brain to the venous blood [37], but it appears that on the other side of the blood-brain barrier (BBB) HSA picks up Aß via intermediates (e.g., lipoproteins) and carries it to liver and kidney, where Aß is cleared [38–40]. The alternative Aß fate is the aggregation into neurotoxic oligomers and then to fibrillogenic species that can ultimately be deposited as Aß amyloid, the hallmark of AD.

Similar to Aß clearance, intracellular Tau can be cleared via Ubiquitin–Proteasome System, Autophagy-Lysosome and proteases, while extracellular Tau can be degraded by enzymes and microglia phagocytosis, or be exported to blood via BBB and blood–cerebrospinal fluid (CSF) barrier [41–43]. Recently, pTau in CSF and periphery blood has become an important AD biomarker [42, 44]. Given the excellent carrier property of HSA and the abundance of this protein, it is reasonable to assume that HSA may also bind with pTau and transport it to liver and kidney for clearance.

### 1.2 Study conclusions and implications

In the current study, we demonstrate that 1) deamidation of HSA leads to its aggregation and thus to a reduction in soluble HSA, which is one of the factors related to clearance of Aß and pTau [33, 34]; 2) deamidation and especially aggregation change HSA structure and reduce the HSA molecules’ ability to bind Aß peptide and pTau protein, which diminishes their clearance; 3) the isoAsp level in HSA as well as the level of free Aß is elevated in AD plasma compared to healthy blood, while 4) the level of anti-aged-HSA antibodies is significantly reduced. Based on the findings above, we updated the isoAsp hypothesis of AD (Figure 1B), which is now formulated as follows: Asn deamidation and subsequent isoAsp formation in HSA lead to its partial unfolding and aggregation, with both processes diminishing the capacity of HSA to carry its ligands, including Aß and pTau. The resulting reduction in clearance enhance Aß and pTau aggregation in brain, increasing the risk of AD. As isoAsp is somewhat immunogenic, there are endogenous antibodies against deamidated HSA that facilitate its removal, and the isoAsp build-up in AD is associated with reduced levels of these antibodies. Altogether, this hypothesis supports the role of deamidation and isoAsp accumulation in AD and offers new possible venues for diagnostics, and possibly prevention, of this debilitating disease.

We also tested whether the obtained LC-MS/MS data contained any evidence of a lower HSA content in AD blood, as literature suggested [33–35], but found no significant difference (P > .05). This could be due to the peculiarity of label-free protein quantification in LC-MS/MS. Although albumin comprises half of all protein content in blood, because of its relative dominance the HSA peptides accounted in LC-MS/MS for > 90% of the total ion current of all identified and quantified peptides, rendering meaningless the conventional normalization by the total peptide abundance. In the absence of proper normalization, the measured HSA abundance was modulated by the total protein concentration in the sample, which fluctuated from sample to sample due to the dilution with preservatives during plasma preparation, masking the differences in the original HSA concentrations. An aggravating factor was also the small size of the reported effect, which was only statistically discernible in a larger cohort [33–35].

Another limitation of our study is the lack of samples from patients with mild cognitive impairment (MCI) and other neurodegenerative diseases, such as Parkinson disease, vascular dementia, etc. Thus, we were unable to correlate the AD progress with the levels of isoAsp in HSA and anti-aHSA antibodies, as well as assess the specificity of the isoAsp-related biomarkers to AD and other neurodegenerative diseases. Also, the predictive power of isoAsp biomarkers needs to be compared with that of other biomarkers used in clinics, such as mental scores, Aß and pTau levels in CSF and blood, sizes of Aß plaques or neurofibrillary tangles, etc. We leave these deficiencies to be filled in future studies.

### 1.4 Discussion

The updated isoAsp hypothesis fits into the current AD etiology narrative (Figure 1B). While the exact cause of AD has not been discovered yet [53, 54], current evidence indicates that brain amyloidosis with or without tauopathy carries the most significant dementia risk, with hazard ratio of 13 at 31% prevalence in the population of cognitively normal persons over 65 years of age [55]. Although hyperphosphorylation of Tau protein occurs after and probably as a result of Aβ plaques deposition [56], the imbalance between the production and clearance of Aß peptides [57] as well as pTau is definitely an aggravating element in AD. Indeed, while AD causes can be multiple, the reduced ability of HSA and other proteins to carry Aß and pTau away from the brain blood barrier has emerged as one of the important factors [25, 28, 29, 35]. One reason for diminished clearance could be the insufficient protein synthesis in the liver, which leads to lower-than-normal HSA concentration in blood. Our modified hypothesis supported by presented experimental data suggests another reason for a bottleneck in Aß and pTau clearance – a failure of repair and/or clearance of deamidated HSA. This failure leads to accumulation of deamidated HSA molecules in blood, with their greatly reduced ability to bind and carry Aß and pTau. As isoAsp repair is unspecific, its failure must affect other ligand carriers as well.

In essence, the main premise suggests that AD may start as a peripheral metabolic disorder that results in increased protein deamidation. An important aggravating factor in this scenario is the reduced levels in blood of anti-aHSA antibodies that help removing deamidated HSA and thus prevent them from forming aggregates with negligible ligand-carrying capacity. The reason why the levels of such antibodies should decrease could be two-fold: on the one hand, the general decline with age of the immune system may lower the antibody production response to protein deamidation, and on the other hand the elevated levels of isoAsp in blood proteins may simply exhaust the supply of such antibodies. These reasons are not mutually exclusive and can act simultaneously.

Moreover, HSA has a number of metal-binding sites that are responsible for metal binding in blood [58]. Ions of metals like copper and iron are found bound to Aβ as well as present at high concentrations in the amyloid plaques, providing support to the metal hypothesis of AD [59, 60]. Also, dysregulated copper ions may initiate and exacerbate tau hyperphosphorylation and formation of pTau fibrils that ultimately contribute to synaptic failure, neuronal death, and cognitive decline observed in AD [61]. A size-exclusion chromatography (SEC) analysis has revealed that HSA is able to decrease the aberrant interaction between Aβ and Cu (II) and revert copper-induced aggregation of Aβ, thus rescuing brain cells from the toxicity of Aß aggregates [62]. It stands to reason that HSA with abnormally high isoAsp content might not have the competence to interrupt the binding of metals with Aβ and pTau.

Besides, inflammation is one of the central mechanisms driving the onset of AD, exacerbating the Aβ and pTau pathology. IsoAsp in blood proteins can trigger inflammation: for example, isoAsp-containing fibronectin activates in mice both monocytes and macrophages, triggering expression of pro-inflammatory cytokines MCP-1 and TNFα to drive additional recruitment of circulating leukocytes [63]. Since an elevated isoAsp level in HSA means deficiency in isoAsp repair or clearance, it also means an elevated risk of inflammation, and constantly elevated isoAsp levels could result in chronic neuroinflammation, reflected in the long term by the reduced levels of anti-isoAsp antibodies in blood.

It is estimated that 79% of dementia cases might be spared by full control of brain amyloidosis and tauopathy [55], which indicates that treatment or prevention of this condition could result in a drastic reduction of AD incidence. The insights provided in this work offer novel ways of diagnosing AD. Since Aß and pTau clearance disruption precedes their aggregation, there is a potential for supplementary early AD diagnostics; this potential needs however be tested experimentally to determine the approach’s sensitivity and specificity.

## 2. CONSOLIDATED RESULTS AND STUDY DESIGN

To test the role of HSA deamidation in AD, we needed an HSA-specific method of isoAsp quantification. LC-MS/MS with electron capture/transfer dissociation (ECD or ETD) MS/MS that has been used in our previous studies [20] possesses the required specificity and would be a method of choice if not for two drawbacks. One drawback is the necessity to employ before the analysis tryptic hydrolysis of proteins, which results in *in vitro* deamidation that could mask the biological differences [45, 46]. The other disadvantage of LC-MS/MS is the low throughput and high analysis cost. The enzyme-linked immunosorbent assay (ELISA) avoids both drawbacks by requiring no protein digestion and being low-cost, high-throughput and scalable. However, such an ELISA required a commercially unavailable antibody specific to deamidated HSA. Thus, we developed a monoclonal antibody to facilitate a high-throughput HSA-specific method of isoAsp quantification [47]. Via the LC-MS/MS analyses of artificially deamidated HSA over time, we discovered an “isoAsp meter” peptide, against which the monoclonal antibody (mAb) 1A3 was raised. An indirect ELISA based on 1A3 mAb provided a proportional response to a mixture of fresh HSA (fHSA) and aged HSA (aHSA) with high sensitivity and specificity (Figure 2A-D).

**Figure 2.**
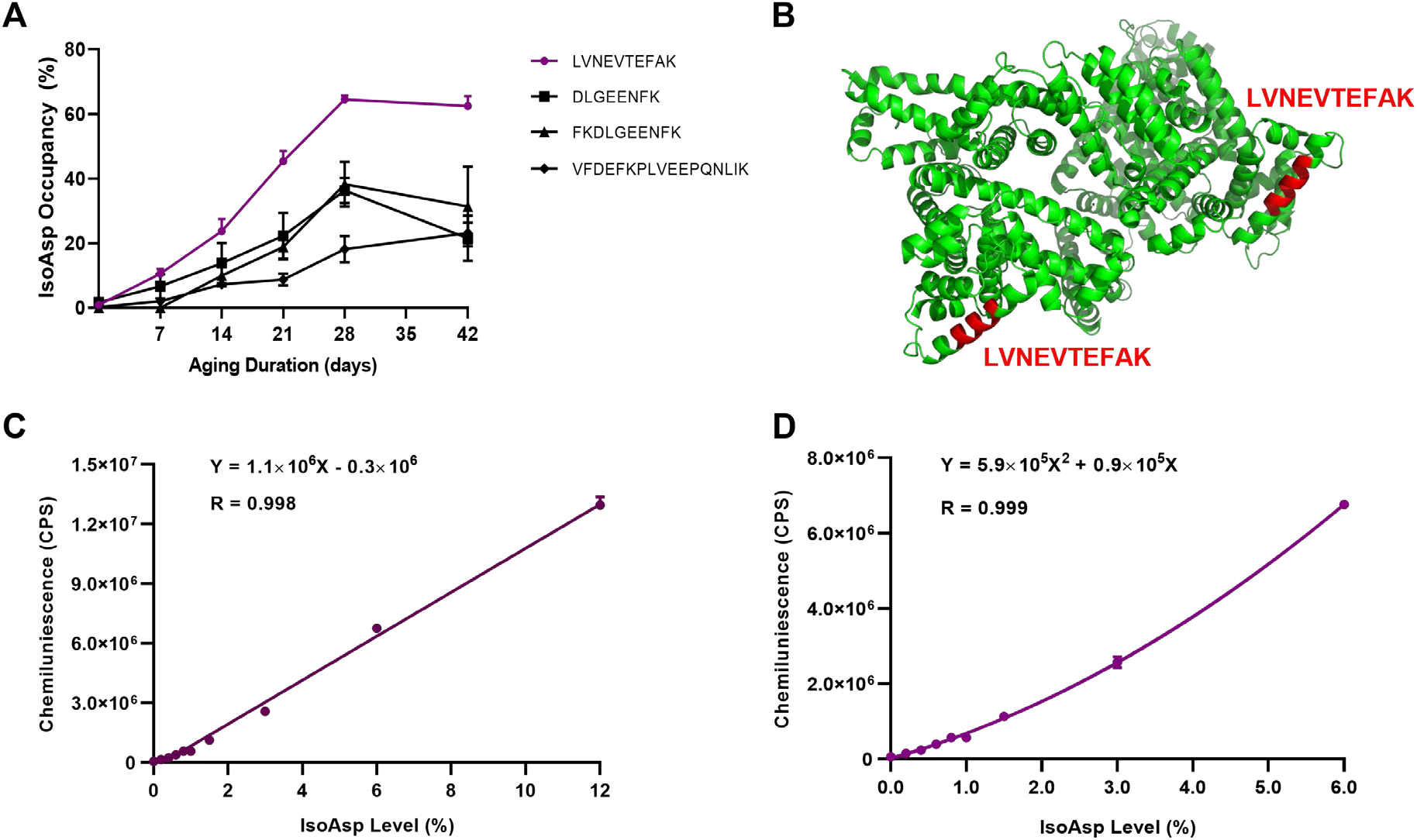
Designing the “IsoAsp Meter” for measuring isoAsp in blood. (A) IsoAsp occupancy changes in different HSA tryptic peptides over aging time. (B) The position of the peptide LVNEVTEFAK in the structure of HSA dimer. (C) ELISA signal from the mixtures of fresh and aged HSA in different proportions up to 12% isoAsp content and a fitted linear calibration curve for isoAsp measurements in blood HSA. (D) Similar for the region of ≤ 6% isoAsp content, where the dependence becomes quadratic. Abbreviations: LC-MS/MS: liquid chromatography tandem mass spectrometry; D*: isoaspartate, isoAsp; D: aspartate, Asp; N: asparagine, Asn; m/z: mass-to-charge ratio.

To further examine the role of isoAsp in AD, we verified by SEC the formation of aggregates in deamidated HSA and a significant increase in AD of the HSA aggregate/monomer ratio (P < 10^-5^) (Figure 3A-G). Furthermore, we found that deamidation significantly diminished the binding capacity of HSA with Aß and pTau (Figure 4A-C), leading to a significantly elevated level of free Aß in AD blood compared to controls (P < .01) (Figure 6E). Molecular dynamic simulations (MDS) of deamidated HSA supported the HSA structural changes upon deamidation (Figure 5A-C). At the same time, we discovered in the same AD cohort significantly lower levels of endogenous antibodies against isoAsp in HSA (P < .0001) and increased levels of isoAsp (P < .001) (Figure 6 A-D), supporting the breakdown in AD of the balance between HSA damage and its repair/clearance.

**Figure 3.**
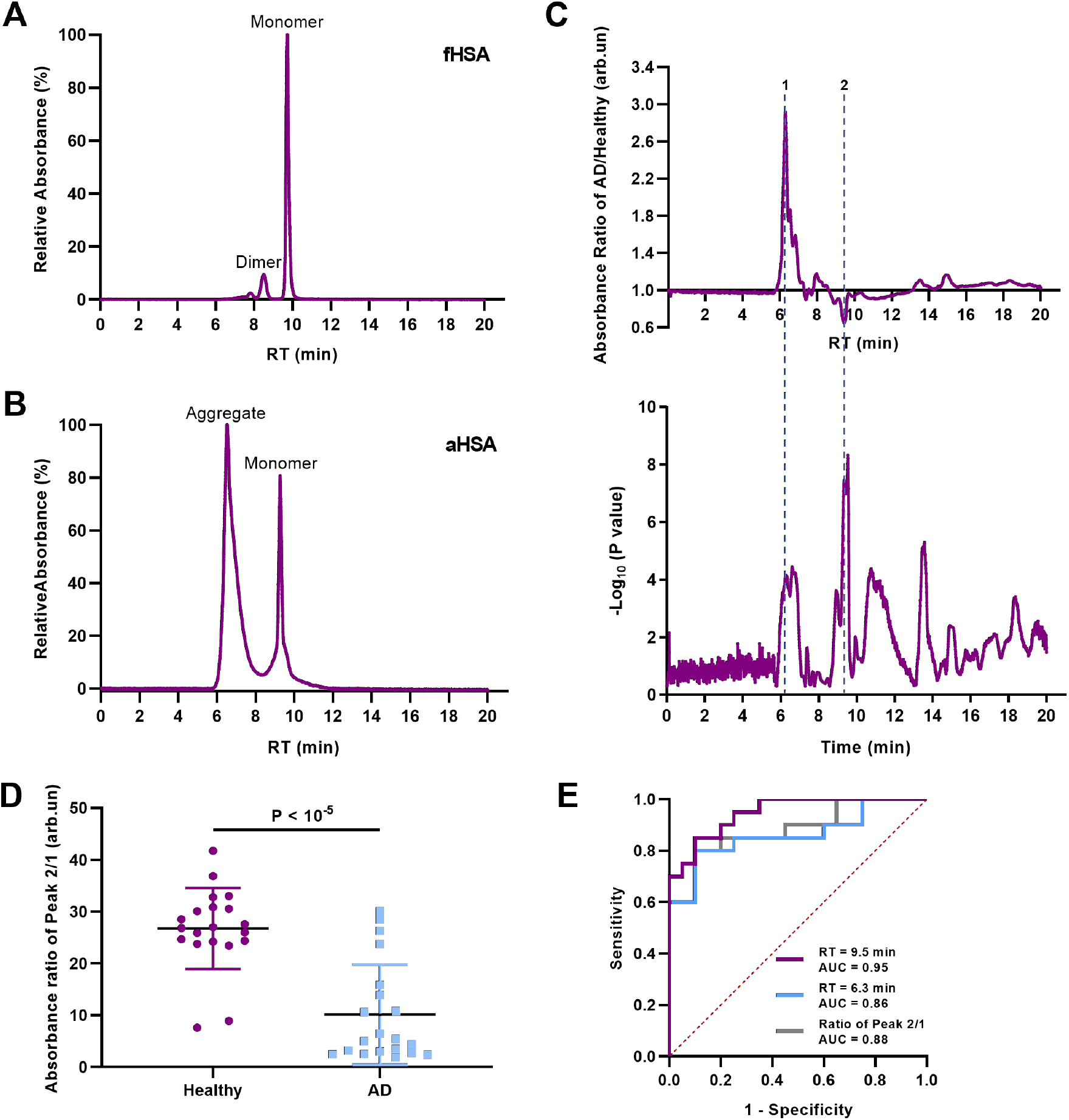
HSA aggregation due to the isoAsp deamidation. (A) SEC shows that HSA monomers comprise more than 90% of fHSA molecules. (B) In aHSA (isoAsp content ≈ 60%), a large portion of aggregates are formed, containing according to SEC ≥ 8 HSA molecules on average. (C) Ratio of the UV absorbance in SEC of the AD and Healthy cohorts (n = 20 + 20) (Up) and the corresponding P values (Down), with the largest differences at RT = 6.3 min (positive Peak 1, P < 10^-4^) and RT = 9.5 min (negative Peak 2, P < 10^-6^). (D) The ratio of the UV absorbance at Peak 2 (RT = 9.5 min) and Peak 1 (RT = 6.3 min). (E) The ROC curves of the comparisons between the AD and Healthy cohorts. Abbreviations: fHSA, fresh human serum albumin; aHSA, aged human serum albumin; Aβ, amyloid-beta; SEC, size exclusion chromatography; RT, retention time; ROC, receiver operating characteristic.

**Figure 4.**
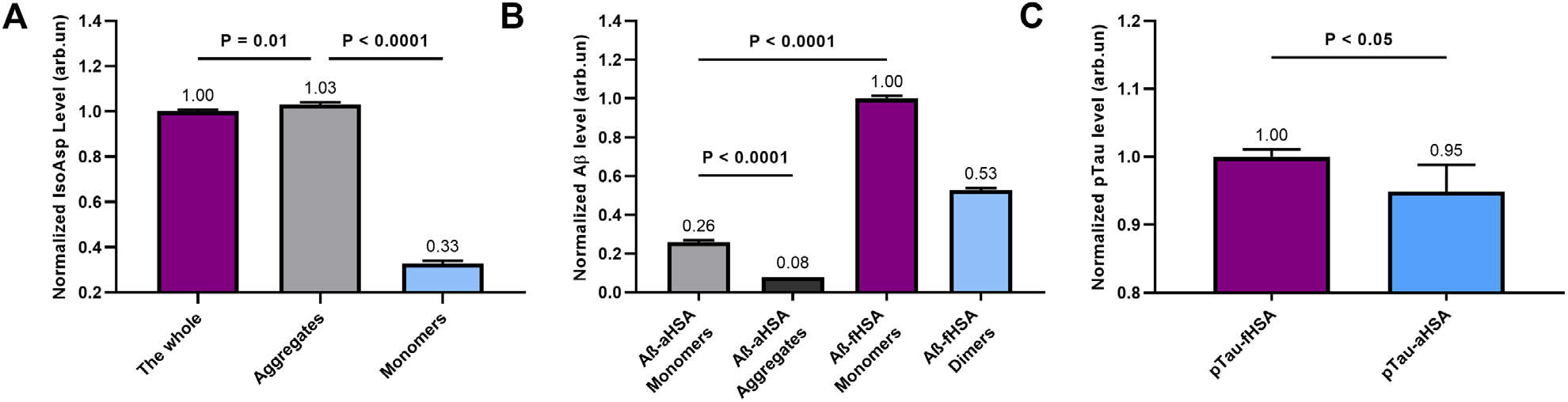
Lower binding of deamidated HSA with Aß and pTau. (A) aHSA aggregates contain 3 times more isoAsp than the aHSA monomers. (B) The aHSA monomers bind 4 times less Aß and aHSA aggregates bind 12 times less Aß than fHSA monomers. (C) The pTau binding by aHSA is significantly lower than that by fHSA. Abbreviations: fHSA, fresh human serum albumin; aHSA, aged human serum albumin; Aβ, amyloid-beta;SEC, size exclusion chromatography; ELISA, enzyme-linked immunosorbent assay.

**Figure 5.**
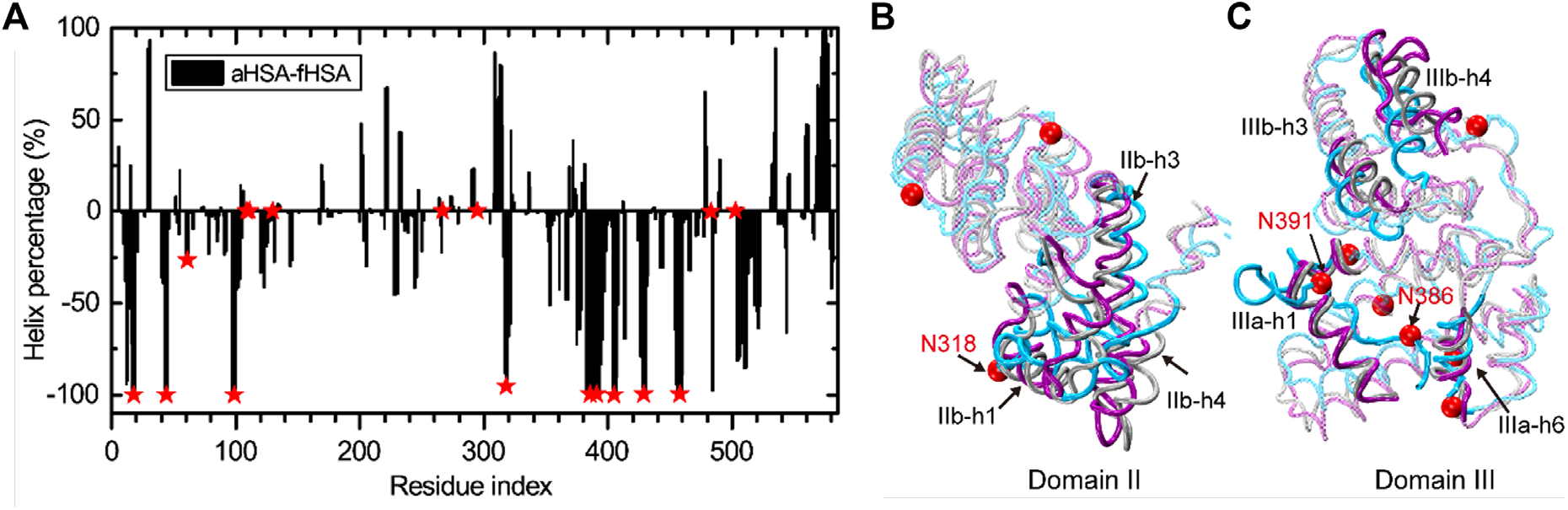
Structural changes of HSA upon deamidation. (A) The difference between the helix percentage in aHSA and fHSA for each residue. Negative values mean the decrease of helix structures in aHSA as compared to fHSA. Deamidation sites are indicated by red stars. (B) and (C) show the superpositions of domain II and III, respectively, in the final state of aHSA (purple) and fHSA (blue) simulations with the crystal structure (gray). Regions with notable structural deviations are opaque while the rest are transparent. Deamidation sites are represented by red spheres. Abbreviations: fHSA, fresh human serum albumin; aHSA, aged human serum albumin.

**Figure 6.**
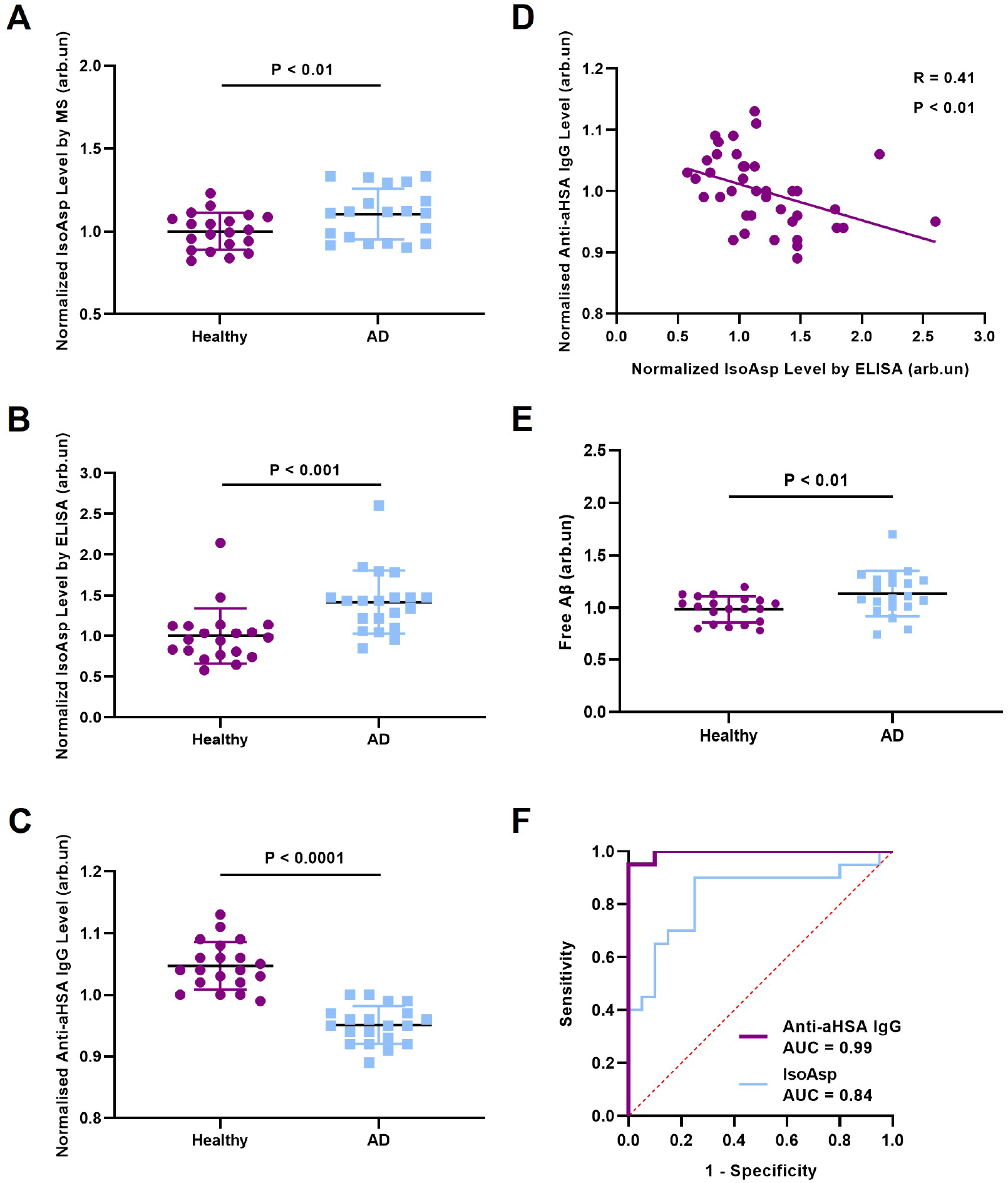
Verification of higher isoAsp content and lower level of anti-isoAsp antibodies in AD blood. (A) The LC-MS/MS analysis reveals that the isoAsp levels in the peptide LVNEVTEFAK are higher in AD patients than healthy controls (n = 20 + 20, P < .01). (B) The isoAsp levels in blood of the same cohort measured by ELISA in three independent experiments show a 30% increase in AD (P < .001). (C) ELISA with aHSA as a bait to capture the IgGs purified from the same cohort shows a 10% higher anti-aHSA IgG level in healthy controls compared to the AD group (P < .0001). (D) The ROC curve on normalized isoAsp levels by LC-MS/MS shows the prediction accuracy of 0.71. (E) An increased level of free Aß in AD blood compared to healthy controls (P < .01). (F) The ROC curve on normalized isoAsp levels by ELISA exhibits the prediction accuracy > 90%. And the AUC generated from the ROC curve of anti-aHSA IgG reaches 0.99. Abbreviations: isoAsp, isoaspartate; mAb, monoclonal antibody; AD, Alzheimer’s disease; HSA, human serum albumin; ELISA, enzyme-linked immunosorbent assay; LC-MS/MS: liquid chromatography tandem mass spectrometry; ROC, receiver operating characteristic; IgG, immunoglobulin G; AUC, area under curve.

## 3. DETAILED METHODS AND RESULTS

### 3.1 Methods

Detailed methods can be found in Supporting Information.

### 3.2 Results

#### 3.2.1 Participant characteristics

Table 1 presents the composition of the AD cohort (n = 20) and the matched healthy cohort (n = 20). No significant differences were observed in age and male percentage between two groups. The Mini-Mental State Examination (MMSE) scores, cerebrospinal fluid (CSF) levels of amyloid β (1–42) (Aβ42), total tau (tTau) and phosphorylated tau (pTau) of AD group are listed. The range of values for CSF biomarkers fell within the measuring ranges of the assays (Aβ42: 430.5–1125.4 pg/mL; tTau: 190.6–847.9 pg/mL; pTau: 20.7–95.9 pg/mL).

**Table 1.** Information of patients from AD group and Healthy group.

| Characteristics | AD (n = 20) | Healthy (n = 20) |
| --- | --- | --- |
| Age, years | 65.5 [60.7, 69.2] | 60.2 [58.9, 61.4] |
| Male, n (%) | 3 (15%) | 8 (40%) |
| MMSE scores | 22.5 [21.1, 25.1] | N/A |
| CSF Aβ42 (pg/mL) | 769.1 [577.3, 925.3] | N/A |
| CSF tTau (pg/mL) | 366.5 [271.0, 423.3] | N/A |
| CSF pTau (pg/mL) | 34.0 [26.7, 43.2] | N/A |

Notes: Age, MMSE scores, CSF Aβ42, CSF tTau and CSF pTau are expressed as median (CI). Sex is expressed as a number of participants and a percentage.
Abbreviations: AD, Alzheimer's disease; MMSE, mini-Mental State Examination; CI, confidence interval; N/A, not available.

#### 3.2.2 Development of the monoclonal antibody against deamidated HSA

After identifying and quantifying the peptides, we compared the isoAsp formation rates in several Asn-containing tryptic peptides (Figure 2A). The peptide LVNEVTEFAK (N = Asn) showed the highest isoAsp occupancy increase, from the background level of 0.9% to ≈ 63%. The final value plateaued after 28 days of deamidation. In the protein structure this peptide is exposed to the solvent, which explains its high deamidation rate and which should facilitate antibody binding to it (Figure 2B).

Since all quantified peptides showed proportional with the ageing time isoAsp occupancy increase, LVNEVTEFAK can be used as an “isoAsp meter” for the whole HSA molecule. The novel monoclonal antibody (mAb) 1A3 raised in a mouse against this peptide is described in detail separately [47]. Here we demonstrate that an indirect ELISA based on 1A3 mAb provides a proportional response to a mixture of fHSA and aHSA (Figure 2C, D), with the signal/noise ratio reaching the threshold value of ≥ 2 already at 0.33% of aHSA admixture, corresponding to 0.2% isoAsp occupancy in LVNEVTEFAK (the average value of isoAsp occupancy in a healthy cohort was found to be 0.74% [47]).

#### 3.2.3 Deamidation causes aggregation of HSA

To verify that deamidation of HSA leads to its aggregation, SEC analysis was performed for fHSA and aHSA. While fHSA gave a major peak corresponding to the HSA monomer, a smaller satellite corresponding to a dimer, and a much smaller oligomer peak (Figure 3A), aHSA produced two strong peaks (Figure 3B), with one peak eluting much earlier than the monomer. The retention time (RT) of that first peak corresponds to the aggregates comprised of eight or more HSA molecules.

Consistent with HSA aggregation with time, we found some of the samples of stored frozen blood plasma to be cloudy, with visible depositions of poorly soluble aggregates.

#### 3.2.4 Distinguishing AD and Healthy blood using SEC

To verify that more HSA aggregates and fewer monomers are present in AD blood, we subjected the plasma samples from AD and Healthy groups (n = 20 + 20, see below) to SEC. The UV absorbance values at a given retention time (RT) were normalized by the total absorbance in each sample and the minimum absorbance in that sample was subtracted. The ratio of the normalized UV absorbances of the two cohorts reached the highest level at Peak 1, RT = 6.3 min (Figure 3C), which corresponds to retention time of the aHSA aggregates (Figure 3B). The absorbance ratio dropped below unity at Peak 2, RT = 9.5 min (Figure 3C), with the retention time corresponding to HSA monomers. The absorbance ratios of Peak 2/1 are shown in Figure 3D. The respective areas under the curve (AUCs) of the receiver operating characteristic (ROC) curves of these data are ranging from 0.86 for 6.3 min absorbance to 0.95 for 9.5 min absorbance (Figure 3E). Since the AUC is usually interpreted as the accuracy of the model separating healthy and disease states [48, 49], there seems to be a potential of plasma SEC analysis for AD diagnostics.

#### 3.2.5 HSA deamidation reduces binding with Aß and pTau

The isoAsp content in the monomers and aggregates of aHSA were analyzed using our 1A3 mAb ELISA [47]. The analysis revealed a ≥ 3 times higher isoAsp content in the aHSA aggregates compared to the monomers (P < .0001, Figure 4A). To verify that deamidated HSA has a reduced ability to bind Aß, we incubated Aß 1-42 with aHSA and fHSA for 2 h at 37 °C and then performed fractionation by SEC. A commercial antibody against Aß 1-42 was then used in indirect ELISA to quantify the amount of Aß in each fraction diluted to the same protein concentration. As expected, fHSA monomers showed the highest binding ability, followed by fHSA dimers, while the aHSA monomers had much lower binding capacity, and aHSA aggregates – negligible binding capacity toward Aß (Figure 4B).

A similar experiment was performed with pTau. We incubated recombinant pTau (MW ≈ 64 kDa) with aHSA and fHSA for 2 h at 37 °C and then isolated the complexes by removing the unbound pTau and HSA monomers by centrifugation through a 100-kDa filter. The complexes of pTau with HSA were quantified by ELISA using anti-pTau monoclonal antibody as the primary antibody. A modest but significant (P < .05) decrease in the aHSA binding capacity with pTau compared to fHSA was found (Figure 4C).

#### 3.2.6 MDS supports reduced capacity of deamidated HSA for ligand binding

To get mechanistic insight into the effect of deamidation on the structure of HSA molecules with MDS, we assumed in a proof-of-concept step that all 17 Asn residues can be deamidated, and simulated the fully deamidated form (denoted as aHSA) in comparison with the wild-type protein (denoted as fHSA). In the 1-microsecond-long simulations, aHSA displayed a larger root-mean-square-deviation (RMSD) from the crystal structure than fHSA (5 vs. 3 Å). As compared to fHSA, the helix probability of aHSA decreased from 0.63 ± 0.01 to 0.53 ± 0.02. The loss of helix structures in aHSA mainly occurred to residues around the deamidation sites (Figure 5A), and the lost helix structures were converted into random coils. Such a structural change increased the aggregation propensity, consistent with the experimentally observed aHSA aggregation.

Next, we used the structure-function relationship to understand the reduced Aß binding capacity of aHSA (pTau binding to HSA is poorly understood and thus was not considered). Since Aß can bind to each of the HSA three domains [50], with the exact binding locations not experimentally proven, we used as plausible binding sites those identified previously in our MDS studies that used two protocols [51, 52]. While for fHSA each of the three domains fitted well the crystal structure (RMSD 2.5 Å), for aHSA only domain I superposed satisfactorily with the crystal structure (RMSD 3.2 Å). Thus, for detailed study we focused on the deviating domains II and III (RMSD 4.3 and 4.4 Å, respectively). In these domains, intriguingly, the regions with prominent structural changes overlapped with the computationally identified Aß binding sites. In domain II (Figure 5B), these are helices h1, h3 and h4 in subdomain IIb, which together with IIb-h2 formed an Aß binding site in our previous simulations. In domain III (Figure 5C), helix h1 in subdomain IIIa and helices h3/h4 in subdomain IIIb with larger structural deviations than the rest of the domain belong to two hotspots for Aß binding. The above results suggest that deamidation of N318 in IIb-h1, N386 and N391 in IIIb-h1 would significantly interfere with Aß binding. Note that IIIb-h3 and IIIb-h4 do not contain Asn residues and their structural changes could be due to intrinsic flexibilities of the C-terminus.

#### 3.2.7 AD blood has elevated isoAsp content in HSA

The first measurement of isoAsp in blood HSA did not reveal any significant difference between the AD and healthy cohorts. However, the verifying analysis performed with LC-MS/MS of digested blood proteins confirmed higher isoAsp levels in the peptide LVNEVTEFAK of AD patients (P = .01; Figure 6A). The tentative explanation of this discrepancy was that the majority of aHSA molecules aggregated, and while in LC-MS/MS analysis the poorly soluble HSA aggregates were mostly digested by trypsin, the isoAsp residues in these aggregates were less amenable to antibody binding in ELISA. To test this hypothesis, we applied 20 min sonication of the blood samples prior to ELISA to break down the aggregates and solubilize the released HSA molecules. The repeated analysis revealed a 30 ± 5% higher isoAsp content in HSA of AD patients compared to healthy controls (P < .001; Figure 6B).

#### 3.2.8 Anti-aHSA antibodies are deficient in AD blood

To test the prediction of deficiency of antibodies against deamidated HSA in AD blood, we purified Immunoglobulin G (IgG) from plasma samples of the same cohorts by Melon Gel IgG Spin Purification Kit. Using aHSA as an antigen, the purified IgGs were employed as primary antibodies in ELISA. The normalized signals from three replicate measurements revealed a 10 ± 1% reduction in anti-aHSA IgGs in AD patients compared to healthy donors (P < .0001) (Figure 6C). There was a negative correlation between the isoAsp content in HSA and anti-aHSA antibody content (Pearson’s R = 0.41, P < .01; Figure 6D). This observation was consistent with the hypothesis that isoAsp in blood HSA combined with a diminished clearance of deamidated HSA could be a risk factor for AD.

#### 3.2.9 Increased levels of Aß not bound with HSA in AD blood

To further confirm the reduced capacity of deamidated HSA binding with Aß, we filtered out proteins larger than 10 kDa, including HSA, and measured in the filtrate of both cohorts the amount of free Aß1-42. There was on average a 10% higher level in AD blood compared to healthy subjects (P < .01; Figure 6E).

#### 3.2.10 Diagnosing AD based on isoAsp-HSA ELISA and anti-aHSA antibody content

For the ROC curve based on the isoAsp-HSA ELISA data calibrated using the second order polymonial regression, the AUC was 0.84. This result suggested potential of blood HSA isoAsp measurements for supplementary AD diagnostics. The AUC for the anti-aHSA antibody ELISA ROC curve was quite impressive, ≈ 0.99 (Figure 6F).

## ACKNOWLEDGEMENTS

This work has been supported by China Scholarship Council (KI-CSC), Horizon 2020 project TopSpec, National Natural Science Foundation of China (Grant 11804218), ZonMw-Memorabel (project no. 73305095003; a project in the context of the Dutch Deltaplan Dementie), Gieskes-Strijbis Foundation, Alzheimer Nederland (Dutch Alzheimer’s Society), Hersenstichting (Dutch Brain Foundation), and Ministry of Science and Higher Education of the Russian Federation (agreement no. 075-15-2020-899). The authors would like to thank Marcus Lundin and Arqum Anwar for their help in the development of the ELISA experiment using mAb against isoAsp in HSA.

## AUTHOR CONTRIBUTIONS

Roman A. Zubarev conceived and supervised the study; Jijing Wang, Roman A. Zubarev and Cong Guo designed research experiments; Charlotte E. Teunissen and Marissa D. Zwan supervised the sample collection in Netherland and provided subject clinical information; Jijing Wang and Zhaowei Meng carried out the SEC experiments; Jijing Wang and Sven Seelow performed the ELISA analyses; Cong Guo and Xin Chen conducted the MDS studies; Susanna L. Lundström and Sergey Rodin provided advice to study design and analysis; Jijing Wang drafted the manuscript; Roman A. Zubarev, Cong Guo and Charlotte E. Teunissen revised the manuscript. All authors read and approved the final manuscript.

## REFERENCES

[1] J.L. Price, J.C. Morris, Tangles and plaques in nondemented aging and “preclinical” Alzheimer’s disease, Ann. Neurol., 45 (1999) 358–368.

[2] J. Cummings, Lessons Learned from Alzheimer Disease: Clinical Trials with Negative Outcomes, Clin. Transl. Sci., 11 (2018) 147–152.

[3] P. Cavazzoni, FDA’s Decision to Approve New Treatment for Alzheimer’s Disease, FDA Center for Drug Evaluation and Research, 2021.

[4] Y.K. Yoo, J. Kim, G. Kim, Y.S. Kim, H.Y. Kim, S. Lee, W.W. Cho, S. Kim, S.M. Lee, B.C. Lee, J.H. Lee, K.S. Hwang, A highly sensitive plasma-based amyloid-beta detection system through medium-changing and noise cancellation system for early diagnosis of the Alzheimer’s disease, Sci. Rep., 7 (2017) 8882.

[5] J.M. Silverman, E. Gibbs, X. Peng, K.M. Martens, C. Balducci, J. Wang, M. Yousefi, C.M. Cowan, G. Lamour, S. Louadi, Y. Ban, J. Robert, S. Stukas, G. Forloni, G.R. Hsiung, S.S. Plotkin, C.L. Wellington, N.R. Cashman, A Rational Structured Epitope Defines a Distinct Subclass of Toxic Amyloid-beta Oligomers, ACS Chem. Neurosci., 9 (2018) 1591–1606.

[6] A. Vojdani, E. Vojdani, Amyloid-Beta 1-42 Cross-Reactive Antibody Prevalent in Human Sera May Contribute to Intraneuronal Deposition of A-Beta-P-42, Int. J. Alzheimers Dis., 2018 (2018) 1672568.

[7] A.L.B. Bitencourt, R.M. Campos, E.N. Cline, W.L. Klein, A. Sebollela, Antibody Fragments as Tools for Elucidating Structure-Toxicity Relationships and for Diagnostic/Therapeutic Targeting of Neurotoxic Amyloid Oligomers, Int. J. Mol. Sci., 21 (2020).

[8] J. Orpiszewski, N. Schormann, B. Kluve-Beckerman, J.J. Liepnieks, M.D. Benson, Protein aging hypothesis of Alzheimer disease, FASEB J., 14 (2000) 1255–1263.

[9] T. Shimizu, A. Watanabe, M. Ogawara, H. Mori, T. Shirasawa, Isoaspartate formation and neurodegeneration in Alzheimer’s disease, Arch. Biochem. Biophys., 381 (2000) 225–234.

[10] B.A. Johnson, J.M. Shirokawa, J.W. Geddes, B.H. Choi, R.C. Kim, D.W. Aswad, Protein L-Isoaspartyl Methyltransferase in Postmortem Brains of Aged Humans, Neurobiol. Aging, 12 (1991) 19–24.

[11] N.E. Robinson, A.B. Robinson, Molecular clocks, Proc Natl Acad Sci U S A, 98 (2001) 944–949.

[12] B. Peters, B.L. Trout, Asparagine deamidation: pH-dependent mechanism from density functional theory, Biochemistry, 45 (2006) 5384–5392.

[13] X. Zhong, J.F. Wright, Biological insights into therapeutic protein modifications throughout trafficking and their biopharmaceutical applications, International journal of cell biology, 2013 (2013).

[14] A.L. Goldberg, Protein degradation and protection against misfolded or damaged proteins, Nature, 426 (2003) 895–899.

[15] L. Bohme, J.W. Bar, T. Hoffmann, S. Manhart, H.H. Ludwig, F. Rosche, H.U. Demuth, Isoaspartate residues dramatically influence substrate recognition and turnover by proteases, Biol. Chem., 389 (2008) 1043–1053.

[16] J.R. Cantor, E.M. Stone, L. Chantranupong, G. Georgiou, The Human Asparaginase-like Protein 1 hASRGL1 Is an Ntn Hydrolase with beta-Aspartyl Peptidase Activity, Biochemistry, 48 (2009) 11026–11031.

[17] T. Noronkoski, I.B. Stoineva, I.P. Ivanov, D.D. Petkov, I. Mononen, Glycosylasparaginase-catalyzed synthesis and hydrolysis of beta-aspartyl peptides, Journal of Biological Chemistry, 273 (1998) 26295–26297.

[18] A.M. Schalk, A. Lavie, Structural and Kinetic Characterization of Guinea Pig L-Asparaginase Type III, Biochemistry, 53 (2014) 2318–2328.

[19] H.A. Doyle, R.J. Gee, M.J. Mamula, Altered immunogenicity of isoaspartate containing proteins, Autoimmunity, 40 (2007) 131–137.

[20] H. Yang, Y. Lyutvinskiy, H. Soininen, R.A. Zubarev, Alzheimer’s disease and mild cognitive impairment are associated with elevated levels of isoaspartyl residues in blood plasma proteins, J Alzheimers Dis, 27 (2011) 113–118.

[21] H.Q. Yang, Y. Lyutvinskiy, S.K. Herukka, H. Soininen, D. Rutishauser, R.A. Zubarev, Prognostic Polypeptide Blood Plasma Biomarkers of Alzheimer’s Disease Progression, Journal of Alzheimers Disease, 40 (2014) 659–666.

[22] M. Cortes-Canteli, D. Zamolodchikov, H.J. Ahn, S. Strickland, E.H. Norris, Fibrinogen and altered hemostasis in Alzheimer’s disease, J Alzheimers Dis, 32 (2012) 599–608.

[23] B.A. Johnson, E.D. Murray, S. Clarke, D.B. Glass, D.W. Aswad, Protein Carboxyl Methyltransferase Facilitates Conversion of Atypical L-Isoaspartyl Peptides to Normal L-Aspartyl Peptides, Journal of Biological Chemistry, 262 (1987) 5622–5629.

[24] X. Hao, Y. Huang, M. Qiu, C. Yin, H. Ren, H. Gan, H. Li, Y. Zhou, J. Xia, W. Li, L. Guo, I.A. Angres, Immunoassay of S-adenosylmethionine and S-adenosylhomocysteine: the methylation index as a biomarker for disease and health status, BMC Res. Notes, 9 (2016) 498.

[25] A.L. Biere, B. Ostaszewski, E.R. Stimson, B.T. Hyman, J.E. Maggio, D.J. Selkoe, Amyloid beta-peptide is transported on lipoproteins and albumin in human plasma, Journal of Biological Chemistry, 271 (1996) 32916–32922.

[26] J. Milojevic, G. Melacini, Stoichiometry and Affinity of the Human Serum Albumin-Alzheimer’s A beta Peptide Interactions, Biophys. J., 100 (2011) 183–192.

[27] M. Zhao, C. Guo, Multipronged Regulatory Functions of Serum Albumin in Early Stages of Amyloid-beta Aggregation, ACS Chem. Neurosci., 12 (2021) 2409–2420.

[28] P. Picon-Pages, J. Bonet, J. Garcia-Garcia, J. Garcia-Buendia, D. Gutierrez, J. Valle, C.E.S. Gomez-Casuso, V. Sidelkivska, A. Alvarez, A. Peralvarez-Marin, A. Suades, X. Fernandez-Busquets, D. Andreu, R. Vicente, B. Oliva, F.J. Munoz, Human Albumin Impairs Amyloid beta-peptide Fibrillation Through its C-terminus: From docking Modeling to Protection Against Neurotoxicity in Alzheimer’s disease, Comput Struct Biotec, 17 (2019) 963–971.

[29] H.F. Stanyon, J.H. Viles, Human serum albumin can regulate amyloid-beta peptide fiber growth in the brain interstitium: Implications for Alzheimer disease, Journal of Biological Chemistry, 287 (2012) 28163–28168.

[30] J. Milojevic, A. Raditsis, G. Melacini, Human Serum Albumin Inhibits A beta Fibrillization through a “Monomer-Competitor” Mechanism, Biophys. J., 97 (2009) 2585–2594.

[31] E.H. Mizrahi, T. Blumstein, M. Arad, A. Adunsky, Serum albumin levels predict cognitive impairment in elderly hip fracture patients, Am J Alzheimers Dis, 23 (2008) 85–90.

[32] D.J. Llewellyn, K.M. Langa, R.P. Friedland, I.A. Lang, Serum Albumin Concentration and Cognitive Impairment, Curr Alzheimer Res, 7 (2010) 91–96.

[33] M. Maes, N. DeVos, A. Wauters, P. Demedts, V. Maurits, H. Neels, E. Bosmans, C. Altamura, A. Lin, C. Song, M. Vandenbroucke, S. Scharpe, Inflammatory markers in younger vs elderly normal volunteers and in patients with Alzheimer’s disease, J. Psychiatr. Res., 33 (1999) 397–405.

[34] T.S. Kim, C.U. Pae, S.J. Yoon, W.Y. Jang, N.J. Lee, J.J. Kim, S.J. Lee, C. Lee, I.H. Paik, C.U. Lee, Decreased plasma antioxidants in patients with Alzheimer’s disease, Int. J. Geriatr. Psychiatry, 21 (2006) 344–348.

[35] J.W. Kim, M.S. Byun, J.H. Lee, D. Yi, S.Y. Jeon, B.K. Sohn, J.Y. Lee, S.A. Shin, Y.K. Kim, K.M. Kang, C.H. Sohn, D.Y. Lee, K.R. Grp, Serum albumin and beta-amyloid deposition in the human brain, Neurology, 95 (2020) E815–E826.

[36] A. Ezra, I. Rabinovich-Nikitin, P. Rabinovich-Toidman, B. Solomon, Multifunctional Effect of Human Serum Albumin Reduces Alzheimer’s Disease Related Pathologies in the 3xTg Mouse Model, J Alzheimers Dis, 50 (2016) 175–188.

[37] M.J. de Leon, K. Blennow, Therapeutic potential for peripheral clearance of misfolded brain proteins, Nature Reviews Neurology, 14 (2018) 637–638.

[38] C.L. Maarouf, J.E. Walker, L.I. Sue, B.N. Dugger, T.G. Beach, G.E. Serrano, Impaired hepatic amyloid-beta degradation in Alzheimer’s disease, Plos One, 13 (2018).

[39] N. Sehgal, A. Gupta, R.K. Valli, S.D. Joshi, J.T. Mills, E. Hamel, P. Khanna, S.C. Jain, S.S. Thakur, V. Ravindranath, Withania somnifera reverses Alzheimer’s disease pathology by enhancing low-density lipoprotein receptor-related protein in liver, Proc Natl Acad Sci U S A, 109 (2012) 3510–3515.

[40] Y.H. Liu, Y.R. Wang, Y. Xiang, H.D. Zhou, B. Giunta, N.B. Manucat-Tan, J. Tan, X.F. Zhou, Y.J. Wang, Clearance of Amyloid-Beta in Alzheimer’s Disease: Shifting the Action Site from Center to Periphery, Mol. Neurobiol., 51 (2015) 1–7.

[41] W.A. Banks, A. Kovac, P. Majerova, K.M. Bullock, M. Shi, J. Zhang, Tau Proteins Cross the Blood-Brain Barrier, J Alzheimers Dis, 55 (2017) 411–419.

[42] S.H. Xin, L. Tan, X. Cao, J.T. Yu, L. Tan, Clearance of Amyloid Beta and Tau in Alzheimer’s Disease: from Mechanisms to Therapy, Neurotox. Res., 34 (2018) 733–748.

[43] Y. Wang, E. Mandelkow, Degradation of tau protein by autophagy and proteasomal pathways, Biochem. Soc. Trans., 40 (2012) 644–652.

[44] B. Olsson, R. Lautner, U. Andreasson, A. Ohrfelt, E. Portelius, M. Bjerke, M. Holtta, C. Rosen, C. Olsson, G. Strobel, E. Wu, K. Dakin, M. Petzold, K. Blennow, H. Zetterberg, CSF and blood biomarkers for the diagnosis of Alzheimer’s disease: a systematic review and meta-analysis, Lancet Neurol., 15 (2016) 673–684.

[45] Y. Du, F.Q. Wang, K. May, W. Xu, H.C. Liu, Determination of Deamidation Artifacts Introduced by Sample Preparation Using O-18-Labeling and Tandem Mass Spectrometry Analysis, Anal. Chem., 84 (2012) 6355–6360.

[46] S.J. Dai, W.Q. Ni, A.N. Patananan, S.G. Clarke, B.L. Karger, Z.H.S. Zhou, Integrated Proteomic Analysis of Major Isoaspartyl-Containing Proteins in the Urine of Wild Type and Protein L-Isoaspartate O-Methyltransferase-Deficient Mice, Anal. Chem., 85 (2013) 2423–2430.

[47] J. Wang, S.L. Lundstrom, S. Seelow, S. Rodin, Z. Meng, J. Astorga-Wells, Q. Jia, R.A. Zubarev, First Immunoassay for Measuring Isoaspartate in Human Serum Albumin, Molecules, 26 (2021).

[48] K. Hajian-Tilaki, Receiver Operating Characteristic (ROC) Curve Analysis for Medical Diagnostic Test Evaluation, Caspian J Intern Med, 4 (2013) 627–635.

[49] J.A. Hanley, B.J. McNeil, The meaning and use of the area under a receiver operating characteristic (ROC) curve, Radiology, 143 (1982) 29–36.

[50] J. Milojevic, G. Melacini, Stoichiometry and affinity of the human serum albumin-Alzheimer’s Abeta peptide interactions, Biophys. J., 100 (2011) 183–192.

[51] H. Xie, C. Guo, Albumin Alters the Conformational Ensemble of Amyloid-beta by Promiscuous Interactions: Implications for Amyloid Inhibition, Front Mol Biosci, 7 (2021) 629520.

[52] C. Guo, H.X. Zhou, Fatty Acids Compete with Abeta in Binding to Serum Albumin by Quenching Its Conformational Flexibility, Biophys. J., 116 (2019) 248–257.

[53] A.A.D.T. Abeysinghe, R.D.U.S. Deshapriya, C. Udawatte, Alzheimer’s disease; a review of the pathophysiological basis and therapeutic interventions, Life Sci., 256 (2020).

[54] L.Y. Fan, C.Y. Mao, X.C. Hu, S. Zhang, Z.H. Yang, Z.W. Hu, H.F. Sun, Y. Fan, Y.L. Dong, J. Yang, C.H. Shi, Y.M. Xu, New Insights Into the Pathogenesis of Alzheimer’s Disease, Front. Neurol., 10 (2020).

[55] G. Livingston, A. Sommerlad, V. Orgeta, S.G. Costafreda, J. Huntley, D. Ames, C. Ballard, S. Banerjee, A. Burns, J. Cohen-Mansfield, C. Cooper, N. Fox, L.N. Gitlin, R. Howard, H.C. Kales, E.B. Larson, K. Ritchie, K. Rockwood, E.L. Sampson, Q. Samus, L.S. Schneider, G. Selbaek, L. Teri, N. Mukadam, Dementia prevention, intervention, and care, Lancet, 390 (2017) 2673–2734.

[56] J.Z. Wang, Y.Y. Xia, I. Grundke-Iqbal, K. Iqbal, Abnormal Hyperphosphorylation of Tau: Sites, Regulation, and Molecular Mechanism of Neurofibrillary Degeneration, Journal of Alzheimers Disease, 33(2013) S123–S139.

[57] D.J. Selkoe, J. Hardy, The amyloid hypothesis of Alzheimer’s disease at 25years, EMBO Mol. Med., 8 (2016) 595–608.

[58] W. Bal, J. Christodoulou, P.J. Sadler, A. Tucker, Multi-metal binding site of serum albumin, J. Inorg. Biochem., 70 (1998) 33–39.

[59] M.A. Lovell, J.D. Robertson, W.J. Teesdale, J.L. Campbell, W.R. Markesbery, Copper, iron and zinc in Alzheimer’s disease senile plaques, J. Neurol. Sci., 158 (1998) 47–52.

[60] A.I. Bush, R.E. Tanzi, Therapeutics for Alzheimer’s disease based on the metal hypothesis, Neurotherapeutics, 5 (2008) 421–432.

[61] K. Zubcic, P.R. Hof, G. Simic, M. Jazvinscak Jembrek, The Role of Copper in Tau-Related Pathology in Alzheimer’s Disease, Front. Mol. Neurosci., 13 (2020) 572308.

[62] L. Perrone, E. Mothes, M. Vignes, A. Mockel, C. Figueroa, M.C. Miquel, M.L. Maddelein, P. Faller, Copper Transfer from Cu-A beta to Human Serum Albumin Inhibits Aggregation, Radical Production and Reduces A beta Toxicity, Chembiochem, 11 (2010) 110–118.

[63] J.E. Park, G. JebaMercy, K. Pazhanchamy, X. Guo, S.C. Ngan, K.C.K. Liou, S.E. Lynn, S.S. Ng, W. Meng, S.C. Lim, M.K. Leow, A.M. Richards, D.J. Pennington, D.P.V. de Kleijn, V. Sorokin, H.H. Ho, N.E. McCarthy, S.K. Sze, Aging-induced isoDGR-modified fibronectin activates monocytic and endothelial cells to promote atherosclerosis, Atherosclerosis, 324 (2021) 58–68.

