## Supplementary information for "Testing the link between isoaspartate and Alzheimer’s disease etiology"

### METHODS

#### Participants

The patients with AD enrolled in the study from the Amsterdam Dementia Cohort Biobank were diagnosed according to the criteria of the National Institute on Aging-Alzheimer's Association (NIA-AA) Research Framework (1). Clinical data was collected through face-to-face interviews by cognitive disorder specialists. Healthy controls at similar ages without cognitive problems were recruited from Dutch Brain Research Registry. Demographic data included age and sex. For AD patients (except two subjects), the mini-mental state examination (MMSE) was scored. And the cerebrospinal fluid (CSF) levels of amyloid β (1– 42) (Aβ42), total tau (t-Tau) and phosphorylated tau (pTau) were measured by the Innotest and Elecsys assays (2). All samples collected under the declaration of Helsinki with informed consent given to the research participants. The information of both AD and healthy groups was blind to people who processed the samples.

#### Discovery of the "IsoAsp Meter" in HSA

*Artificial Deamidation of Human Serum Albumin (HSA)*

To imitate natural aging process in a short time, three tubes (1 mL in each tube) of 5 mg/mL HSA (Sigma Aldrich) were incubated in Tris pH 8.5 buffer at 60 ˚C for 7, 14, 21, 28 and 42 days respectively. All samples were stored in -80 ˚C for further preparation and analysis.

#### Quantification of IsoAsp Occupancy in HSA

*Sample Preparation*

A tube of totally fresh HSA (fHSA) was also prepared in the same buffer, and all HSA samples (both fHSA and deamidated ones) were diluted in 8 M Urea Tris pH 8.5 buffer to 1 mg/mL. Each sample containing 5 µg HSA was reduced with 20 mM dithiothreitol for 60 min at 37 °C and alkylated with 66 mM iodoacetamide for 30 min in the dark. Trypsin was added at 1:50 ratio (enzyme: protein, w/w) for digestion at 37 °C for 6 h. Tryptic peptides were desalted by Sep Pak C18 cartridges (Waters), dried in the speed vac and resuspended in 0.1% formic acid and 1% acetonitrile.

*Mass Spectrometry Analysis*

A nano-liquid chromatography (LC) system Ultimate 3000 connected in-line to a Fusion Orbitrap mass spectrometer (both – Thermo Fisher Scientific) was used. Reversed phase LC-separation of the peptides (Gradient A: water with 2% acetonitrile and 0.1% formic acid and Gradient B: acetonitrile with 2% water and 0.1% formic acid) was performed on a 50 cm long EASY spray column (PepMap, C18, 3µm, 100Å) for 240 min per sample. The gradient was set up as follows: 5% B for 5 min, 1–30% B in 95 min, 31–95% B in 5 min, 95% B for 5 min and 1% (B) for 10min. The flow rate was set at 300 nL/min. Precursor ion fragmentation was performed with simultaneous higher-energy collision dissociation (HCD; collision energy: 27%, resolution 15,000, AGC target 5.0e4, maximum injection time 200 ms) and electron transfer dissociation (ETD; "collision energy": 40%, resolution 15,000, AGC target 5.0e4, maximum injection time 200 ms).

*IsoAsp Identification*

MS/MS spectra were extracted using a home-written program RAW to MGF and the resultant .mgf files were searched on Mascot to get .dat files. The data were processed on another in-house software Quanti (3) to quantify the isoAsp occupancy in the Asn-containing peptides over the aging duration. The peptide sequence with the highest rate of isoAsp occupancy increase was chosen as the "IsoAsp Meter".

**Development of a monoclonal antibody-based assay to quantify isoAsp in HSA**

*Development of Monoclonal Antibodies (mAbs) against IsoAsp*

The variants of "IsoAsp Meter" with Asn, Asp and isoAsp were synthesized by SynPeptide (Shanghai, China). The hybridoma clones from mice expressing mAbs targeting the isoAsp peptide but not to Asn or Asp peptide were produced by the Russian Research Center of Molecular Diagnostics and Therapy. The clone 1A3 with highest specificity and signal to noise ratio were selected for following studies.

*Determination of isoAsp in aHSA by Indirect Enzyme-Linked Immunosorbent Assay (ELISA)*

The aHSA samples were prepared in four replicates as 5 mg/mL in PBS buffer. After separation via SEC as above, we collected the fractions of aHSA monomer and aggregates, and their concentrations were analyzed by BCA Protein Assay Kit (Thermo Fisher Scientific). Then the isoAsp content in the monomers and aggregates of aHSA were analyzed by the isoAsp-HSA ELISA as described before (4).

**Size exclusion chromatography (SEC) analysis of fHSA, aged HSA (aHSA) and plasma samples**

The SEC-UV analysis was performed on a Thermo Fisher UltiMate 3000 RSLC-system (Thermo Scientific, Dionex Softron GmbH) equipped with VWD. The separation was in isocratic mode with 0.3 mL/min 1X PBS (HyClone) in an ACQUITY UPLC Protein BEH SEC column (200Å, 1.7 µm, 4.6 mm x 300 mm; Waters). The total chromatographic run time was 20 min. Data acquisition and integration of the chromatographic peaks were controlled by Chromeleon software (Thermo Scientific). Absorption was recorded at wavelength of 214, 254 and 280 nm. The fHSA and aHSA samples were prepared at concentration of 5 mg/mL in PBS, while the plasma samples were diluted to 20 mg/mL in PBS.

**Molecular dynamics simulations of aHSA and fSA**

Protein molecules were described by amber99sb force field (5). According to a force field assessment for β-amino acids (6), the amber99sb force field showed the best accuracy against NMR data. The force field of the IsoAsp residue was taken from the same study. The initial structure of fHSA was taken from the crystal structure of HSA (PDB 1AO6) (7). The initial structure of aHSA was built upon fHSA by replacing all 17 Asn residues with IsoAsp. The chemical structure of IsoAsp was taken from the crystal structure of the deamidated lysozyme protein (PDB 1AT6) . Each Asn residue was manually replaced with the IsoAsp101 residue in PDB 1AT6 after structural superposition. The local structural clashes were removed by subsequent energy minimization. For both systems, protein was solvated in an octahedron box with TIP3P water. After energy minimization and equilibration with protein heavy atoms restrained, a single production run at 310 K and 1 bar was performed for each system. Long-range electrostatic interactions were calculated with the particle mesh Ewald method. Van der Waals interactions were calculated using a cutoff of 1 nm. The simulation time is 1 microsecond. The last 200 ns was used for analysis. All simulations and analysis were carried out with GROMACS-2019 software package (8).

**Measurement of the binding capacity of aHSA and fHSA with amyloid beta (Aβ) and phosphorylated Tau (pTau)**

For testing the binding capacity of aHSA with Aβ, 50 µL of 100 µM Aβ 1-42 (Abcam) was incubated with 50 µL 100 µM aHSA and fHSA respectively for 2 h at 37 °C and then fractionation was performed by SEC. The anti-Aß 1-42 mAb (Sigma Aldrich) was then used in indirect ELISA to quantify the amount of Aß in each fraction diluted to the same protein concentration. A similar experiment was performed with pTau. The recombinant pTau (Sigma Aldrich) with aHSA and fHSA for 2 h at 37 °C and then the antigen-antibody complexes were purified via the 100-kD Amicon Ultra Centrifugal filters (Merck). The complexes of pTau with HSA were diluted to 5 ug/mL and then quantified by indirect ELISA using monoclonal antibody against pTau at Ser-396 (Thermo Fisher Scientific) as the primary antibody.

**Determination of isoAsp in blood HSA**

The plasma samples from AD patients (n = 20) and healthy controls (n = 20) were sonicated in the ultrasonic bath (VWR, Philadelphia, PA, USA) for 20 min and centrifuged at 20,000 × g (Eppendorf, San Diego, CA, USA) for 10 min. For each sample, the supernatant was collected, and the protein concentration was measured by BCA Protein Assay Kit (Thermo Fisher Scientific). All samples were analyzed in triplicates via the same indirect ELISA for aHSA as described above (4). The isoAsp occupancy in each blood sample was calculated according to the standard curve of HSA from each plate. The statistical significance was analyzed by unpaired two tailed t test that yielded p-values.

**Analysis of Aβ not bound with HSA in blood**

100 µL of each centrifuged plasma sample from above was transferred to the 10 kDa Amicon Ultra-0.5 Centrifugal Filter (Sigma Aldrich). After centrifugation at 14,000 x rpm for 30 min, collected the filtered solution in the tube and topped up to 100 µL using the samples dilution buffer from the Human Aβ Ultrasensitive ELISA kit (Thermo Fisher Scientific). Prepared the Aβ standards and conducted the sandwich ELISA according to the manufacturer's manual.

**Determination of anti-aHSA antibody levels in blood**

*Melon Gel Purification*

The IgGs were purified from plasma samples by the Melon Gel IgG Purification Kit (Thermo Fisher Scientific) according to the manufacturer's protocol. The protein concentration was measured by BCA Protein Assay Kit and the purified IgGs were stored at -80 ˚C until further uses.

*Indirect ELISA on Anti-aHSA Antibodies*

The 96-well opaque white polysterene plates were incubated with 50 µL of 5 µg/mL aHSA (≈60% isoAsp) at room temperature for 2 h. Plates were rinsed thrice between steps with PBST. After incubating with blocking buffer at room temperature for 2 h, 50 µL purified IgG from each plasma sample (diluted to 800 ng/mL in blocking buffer) was added respectively at 4 °C for overnight, followed by incubating with 50 µL secondary antibody - Goat anti-Human IgG (H+L) Secondary Antibody conjugated with HRP (Thermo Fisher Scientific) at room temperature for 2 h. The chemiluminescence was measured immediately after adding the working substrates as above. The data were normalized via dividing the intensity of each sample by the average intensity of each row and column.

**Statistics Analyses**

All data of the AD group were normalized via dividing by the median values of Healthy group. All plots were processed by GraphPad Prism (version 8.0.2). Mann-Whitney tests were performed to calculate P values between two groups.

### REFERENCES


1. Jack CR, Jr., Bennett DA, Blennow K, Carrillo MC, Dunn B, Haeberlein SB, et al. NIA-AA Research Framework: Toward a biological definition of Alzheimer's disease. Alzheimers Dement. 2018;14(4):535-62.
2. Willemse EAJ, van Maurik IS, Tijms BM, Bouwman FH, Franke A, Hubeek I, et al. Diagnostic performance of Elecsys immunoassays for cerebrospinal fluid Alzheimer's disease biomarkers in a nonacademic, multicenter memory clinic cohort: The ABIDE project. Alzheimers Dement (Amst). 2018;10:563-72.
3. Lyutvinskiy Y, Yang H, Rutishauser D, Zubarev RA. In silico instrumental response correction improves precision of label-free proteomics and accuracy of proteomics-based predictive models. Mol Cell Proteomics. 2013;12(8):2324-31.

4. Wang J, Lundstrom SL, Seelow S, Rodin S, Meng Z, Astorga-Wells J, et al. First Immunoassay for Measuring Isoaspartate in Human Serum Albumin. Molecules. 2021;26(21).
5. Hornak V, Abel R, Okur A, Strockbine B, Roitberg A, Simmerling C. Comparison of multiple Amber force fields and development of improved protein backbone parameters. Proteins. 2006;65(3):712-25.
6. Paissoni C, Nardelli F, Zanella S, Curnis F, Belvisi L, Musco G, et al. A critical assessment of force field accuracy against NMR data for cyclic peptides containing beta-amino acids. Phys Chem Chem Phys. 2018;20(23):15807-16.
7. Sugio S, Kashima A, Mochizuki S, Noda M, Kobayashi K. Crystal structure of human serum albumin at 2.5 angstrom resolution. Protein Eng. 1999;12(6):439-46.
8. Noguchi S, Miyawaki K, Satow Y. Succinimide and isoaspartate residues in the crystal structures of hen egg-white lysozyme complexed with tri-N-acetylchitotriose. J Mol Biol. 1998;278(1):231-8.
